# Comparison of genotoxic vs. non-genotoxic stabilization of p53 provides insight into parallel stress-responsive transcriptional networks

**DOI:** 10.1101/360974

**Authors:** Allison N. Catizone, Shelley L. Berger, Morgan A. Sammons

## Abstract

The tumor suppressor protein p53 is activated in response to diverse intrinsic and extrinsic cellular stresses and controls a broad cell-protective gene network. Whether p53:DNA binding and subsequent transcriptional activation differs downstream of these diverse intrinsic and extrinsic activators within the same cell type is controversial. Using primary human fibroblasts, we assessed the genome-wide profile of p53 binding, chromatin structure, and transcriptional dynamics after either genotoxic or non-genotoxic activation of p53. Activation of p53 by treatment with either etoposide or the small molecule MDM2 inhibitor nutlin 3A yields strikingly similar genome-wide binding of p53 and concomitant changes to local chromatin modifications and structure. DNA damage, but not p53 activation *per se*, leads to increased expression of genes in an inflammatory cytokine pathway. Etoposide-mediated activation of this inflammation signature is inhibited by treatment with the NF-kB pathway inhibitor Bay 11-7082, but does not affect expression of canonical p53 target genes. Our data demonstrate that differential activation of p53 within the same cell type leads to highly similar genome-wide binding, chromatin dynamics, and gene expression dynamics, and that DNA damage-mediated signaling through NF-κB likely controls the observed pro-inflammatory cytokine gene expression pattern.

## Introduction

The transcription factor p53 serves as a central hub in the transcriptional response to DNA damage(1). p53 directly binds a consensus response element (RE) sequence within gene promoters and enhancers to activate a cell and organism-protective gene regulatory network (2). This transcriptional response involves upregulation of numerous genes involved in cell cycle arrest, apoptosis, DNA repair, and metabolic pathways (3, 4). As loss of p53 activity is highly correlated with tumorigenesis (1), there is strong and continued interest in deciphering the gene networks downstream of wild-type p53 activation.

The p53 protein is normally kept inactive through proteosome-dependent degradation mediated by the E3 ubiquitin ligase MDM2(5). Upon the onset of DNA damage, the ATM and ATR kinases signal through CHK1 and CHK2 to phosphorylate p53, thus liberating active p53 from MDM2-mediated ubiquitination and turnover (6). Nutlin 3A is an MDM2 antagonist that leads to rapid stabilization and activation of p53 protein in the absence of DNA damage and ATM/ATR signaling (7). Importantly, nutlin 3A is highly specific for the p53:MDM2 interaction with transcriptional profiling demonstrating essentially no off-target gene expression changes after treatment (8). Chemical derivatives of nutlin 3A are still under investigation for the treatment of wild-type p53-containing cancers due to the high specificity and seemingly non-genotoxic mechanism of action (9, 10). Nutlin 3A, along with other non-genotoxic small molecule p53 activators, has become a highly used laboratory tool for p53 stabilization without affecting potential parallel DNA damage pathways.

The dynamics of p53 protein stabilization and subsequent cellular-level phenotypes depend on the method used to activate p53, with significant differences observed within different DNA damage paradigms or nutlin 3A (11, 12). Exposure to gamma irradiation lead to oscillating p53 protein levels over a 24-hour period whereas UV treatment produces sustained p53 levels with an overall higher amplitude. In contrast, single doses of nutlin 3A lead to rapid p53 stabilization that is later reversed due to both nutlin 3A degradation and increased p53-dependent expression of MDM2. These p53 dynamics appear to control the ultimate outcomes of p53 activation such as the decision to commit to senescence or apoptosis, for example (11, 12). Although the dynamics of p53 protein levels are directly influenced by the method of p53 stabilization, whether this leads to differential p53:DNA binding or gene activation is less clear (13–16).

The first wave of genome-scale p53 ChIP-seq experiments suggested high spatial variability of p53 binding in response to various p53 activating conditions, even within the same cell type (14, 15, 17–19). Reanalysis of these datasets and multiple other p53 ChIP-seq datasets from a variety of transformed cell types and p53 stabilizaion methods suggested p53 DNA binding is much less variable (13, 20). Approximately 1,000 p53 binding sites display high concordance across multiple labs, cell types, and experimental methods when consistent data processing methods are used (13). Conversely, two recent pre-prints demonstrate widespread cell type-specific p53 binding events that are driven by differences in chromatin accessibility (21, 22). A recent multi-omics approach suggests that high affinity p53 binding sites are shared across cell types, whereas the observed cell type-specific binding events were lower affinity sites (23).

We therefore sought to better understand functional differences between genotoxic and non-genotoxic stabilization of p53 and the resulting transcriptomes. Here, we find that stabilization of p53 by genotoxic (etoposide) and non-genotoxic (nutlin 3A) methods yield nearly identical DNA binding within highly similar local chromatin environments. Direct p53 binding sites are characterized by high levels of H4K16ac, while indirect ChIP-seq-derived p53 binding events are found within highly accessible, promoter regions. Genotoxic activation of p53 using etoposide leads to significantly more activated gene targets than using nutlin 3A, with the majority of these genes classified as inflammatory response genes. Expression of these etoposide-activated genes is abrogated by treatment with NF-κB pathway inhibitors, suggesting a DNA damage-dependent, but p53-independent, mechanism of action. These data provide increased evidence that p53 engagement with the genome and transcriptional targets are cell type-intrinsic and that careful analysis of crosstalk between DNA damage signaling pathways is prudent.

## Results

### Comparison of p53 interaction with the genome after genotoxic and non-genotoxic activation

We used low-passage (PD 25-30) primary human fibroblasts (IMR90) cultured under normoxic conditions (3% O_2_) to assess whether p53-mediated gene expression and genome binding dynamics vary based on the method of p53 stabilization and activation. Of note, the majority of published datasets regarding p53 activity have been performed using 20% O_2_. Etoposide is a commonly used chemotherapeutic that inhibits topoisomerase II, leading to a failure to resolve dsDNA breaks and activation of p53 through an ATM-mediated signaling cascade (24, 25). Phosphorylation of p53 at serine 15 disrupts the interaction with the E3 ligase MDM2 and results in stabilization of the p53 protein (26). The small molecule Nutlin-3A is an inhibitor of the p53:MDM2 interaction, and leads to stabilization and activation of p53a in the absence of DNA damage or p53-S15ph (Fig.1A, (7) Treatment with either 100uM Etoposide or 5uM nutlin 3A lead to similar p53 and p21 (a canonical target of p53) protein accumulation 6 hours post-treatment compared to a DMSO vehicle control (Fig. 1A), suggesting approximately equivalent effects on p53 stabilization and activity. Etoposide treatment led to an increase in phosphorylation of serine 15 (Fig. 1A), which is downstream of DNA damage-dependent kinases and is required for endogenous stabilization of p53 after DNA damage (1).

**Figure 1.**
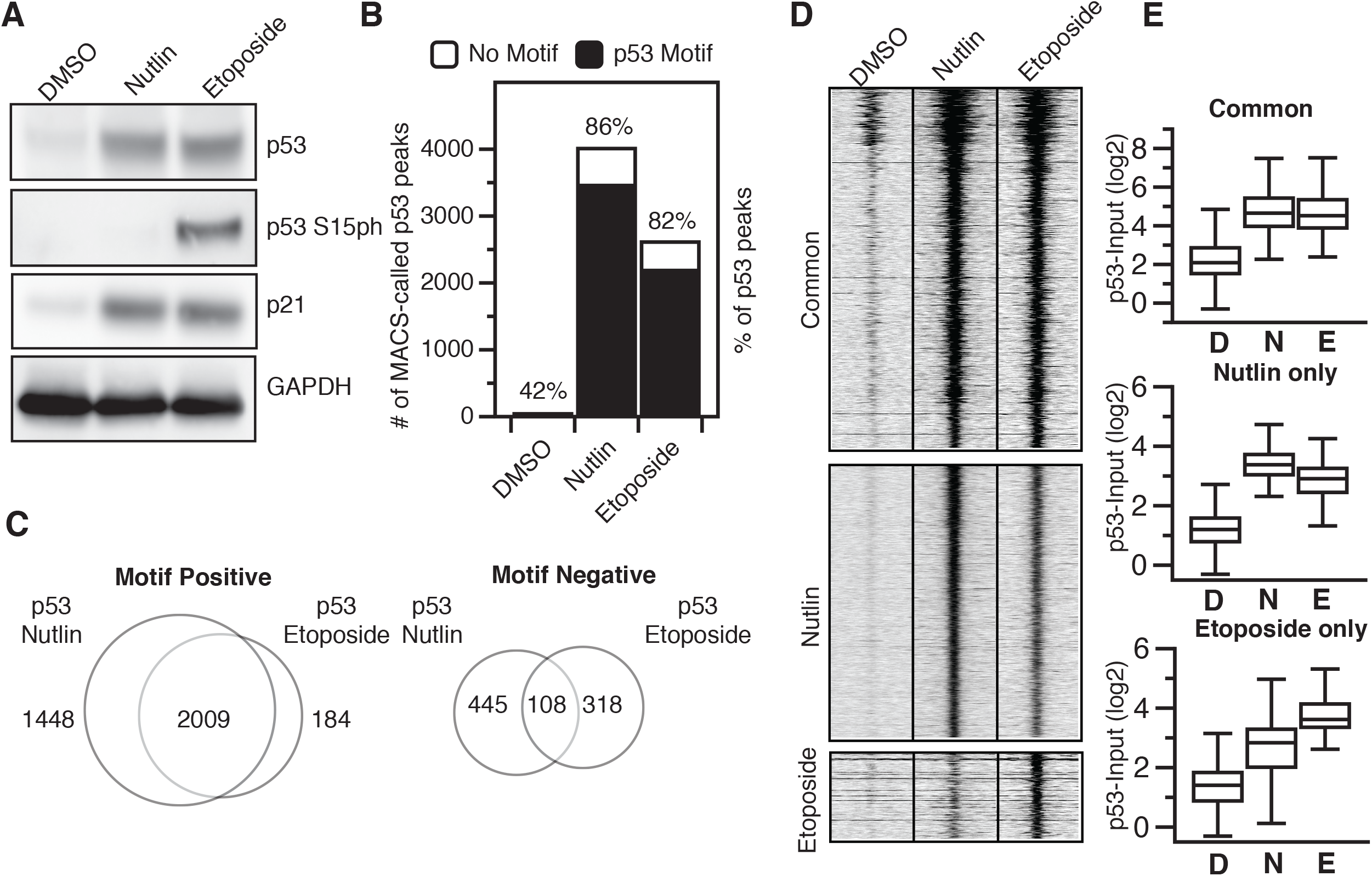
*A*. Western blot analysis of p53, p53 serine 15 phosphorylation, p21, and GAPDH in IMR90 fetal lung fibroblasts (cultured at 3% O_2_ in DMEM plus 10% fetal bovine serum) six hours after treatment with either DMSO (vehicle), nutlin 3A (5uM final), or etoposide (100uM final). *B*. The number of MACS (v1.4) derived p53 peaks (q<0.05) with (black) or without (white) a canonical p53-motif (as determined by p53scan) under DMSO, nutlin 3A, or etoposide treatment conditions. *C*. Intersection between nutlin 3A and etoposide p53 peaks containing a p53 motif (left) or lacking a p53 motif (right) as determined by BedTools intersectBed. *D*. Input-subtracted p53 ChIP-seq tag enrichment at common, nutlin 3A-specific, or etoposide-specific p53-motif containing peaks (-/+ 1000bp from p53 motif center). *E*. Box plot quantification showing input-subtracted p53 enrichment (log2, -/+ 1000bp) at common (top), nutlin 3A-specific (middle), or etoposide-specific (bottom) p53 motif-containing peaks.

We then used chromatin immunoprecipitation coupled to highly parallel sequencing (ChIP-seq) to determine the genomewide binding sites of p53 after 6 hours of treatment with 100uM etoposide and compared this treatment to previously published datasets for DMSO and nutlin (27). Importantly, all p53 ChIP-seq experiments were performed using identical conditions (27). The resulting genome alignment and peak calling data for all experiments can be found in Supplemental Table 1. Treatment with either nutlin or etoposide dramatically increased the number of observed input-normalized p53 peaks compared to DMSO vehicle controls (Fig. 1B), with more statistically enriched peaks (FDR > 0.01) observed after treatment with nutlin 3A than with etoposide. The large majority of p53 binding events in both conditions contain a full canonical p53 response element motif (86% and 82% for nutlin 3A and etoposide, respectively) as determined by p53scan (19), and *de novo* motif finding using HOMER (28) yielded highly similar DNA elements underlying nutlin 3A and etoposide induced p53 binding sites (Supplemental Table 2).

In order to identify putative functional differences between two p53 activating conditions, we analyzed whether nutlin and etoposide-induced p53 binding events occurred within similar genomic loci. We parsed peaks by the presence of a canonical p53 response element motif (motif positive) and those lacking such a motif (motif negative) using p53scan (19) and then performed peak overlap analysis using bedTools (29). Over 90% of etoposide p53 motif+ peaks intersect with nutlin 3A motif+ peaks (Fig. 1C, left), while we observe nearly 1,500 nutlin 3A-specific p53 binding events. Conversely, only 25% of etoposide p53 peaks lacking a canonical p53 motif overlap motif-peaks found after nutlin 3A treatment (Fig. 1C, right). These peak-based results are similar to previous reports of p53 binding after stabilization using various p53 activation paradigms (14,15).

We next examined the ChIP enrichment of motif-positive common, nutlin 3A-specific, and etoposide-specific p53 binding events to determine more quantitative differences between the groups. Despite the seemingly large number of observed nutlin 3A-specific p53 enriched peaks relative to etoposide (Fig. 1C, left), the enrichment of p53 signal within each peak region is well correlated between nutlin and etoposide treatments (Figs. 1D-E). The same is true when looking at enrichment of nutlin-induced p53 binding events at supposedly etoposide-specific locations (Figs. 1D-E). Overall, enrichment at nutlin 3A and etoposide p53 binding events with p53 motifs are well correlated (Pearson ⍴=0.9451), while enrichment of peaks lacking p53 motifs are uncorrelated (Pearson ⍴=0.0134). These data suggest that virtually all inducible p53 binding events are observed independent of p53 activation method when considering enrichment instead of strict peak calling methods. This is in contrast to previous reports using peak calling methodologies (14, 15), but similar to meta-analyses of those (and other) data showing high similarity across p53 conditions when considering ChIP enrichment (13). Our data demonstrate that p53 engagement with the genome is highly similar between non-genotoxic (nutlin 3A) and genotoxic (etoposide) stabilization methods. Further, these results suggest that p53 ChIP-seq peaks lacking canonical p53 motifs are quite variable and do not correlate between stabilization conditions, suggesting they represent experimental or technical artifacts commonly observed in ChIP-seq experiments.

### Chromatin context at p53 binding sites provides evidence for common gene regulation downstream of nutlin 3A and etoposide-mediated activation of p53

p53 ChIP-seq peaks containing a canonical p53 motif (motif +) are located significantly further from transcriptional start sites (TSS) than peaks lacking a canonical motif (motif -), with the modal group of motif – peaks located within 5kb of a TSS (Fig. 2A). Transcriptional start sites and highly expressed genes can cause significant artifacts and false-positives in ChIP-seq experiments (30, 31). We therefore sought to better understand both groups of p53 ChIP-seq peaks by extending our analysis to include chromatin context at p53 binding sites. Specific chromatin structure and modifications are associated with transcriptional regulatory regions, such as the high enrichment of H3K4me3 at promoters/TSS and low H3K4me3/high H3K27ac at enhancer regions (32). p53 binding occurs predominantly within cis-regulatory regions, like enhancers and promoters, in primary skin fibroblasts (27, 33). Thus, we compared p53 binding locations with regions of enriched enhancer and promoter-associated chromatin modifications. Global histone modification levels for transcriptionally-associated H3K4me1/2/3, H3K27ac, and H4K16ac were highly similar across treatment conditions as determined by western blotting (Fig. 2B). We then performed ChIP-seq for these modifications (and total histone H3) under etoposide-treated conditions to determine how DNA damage-associated chromatin dynamics compare to previous observations after DMSO and nutlin-3A treatment (27).

**Figure 2.**
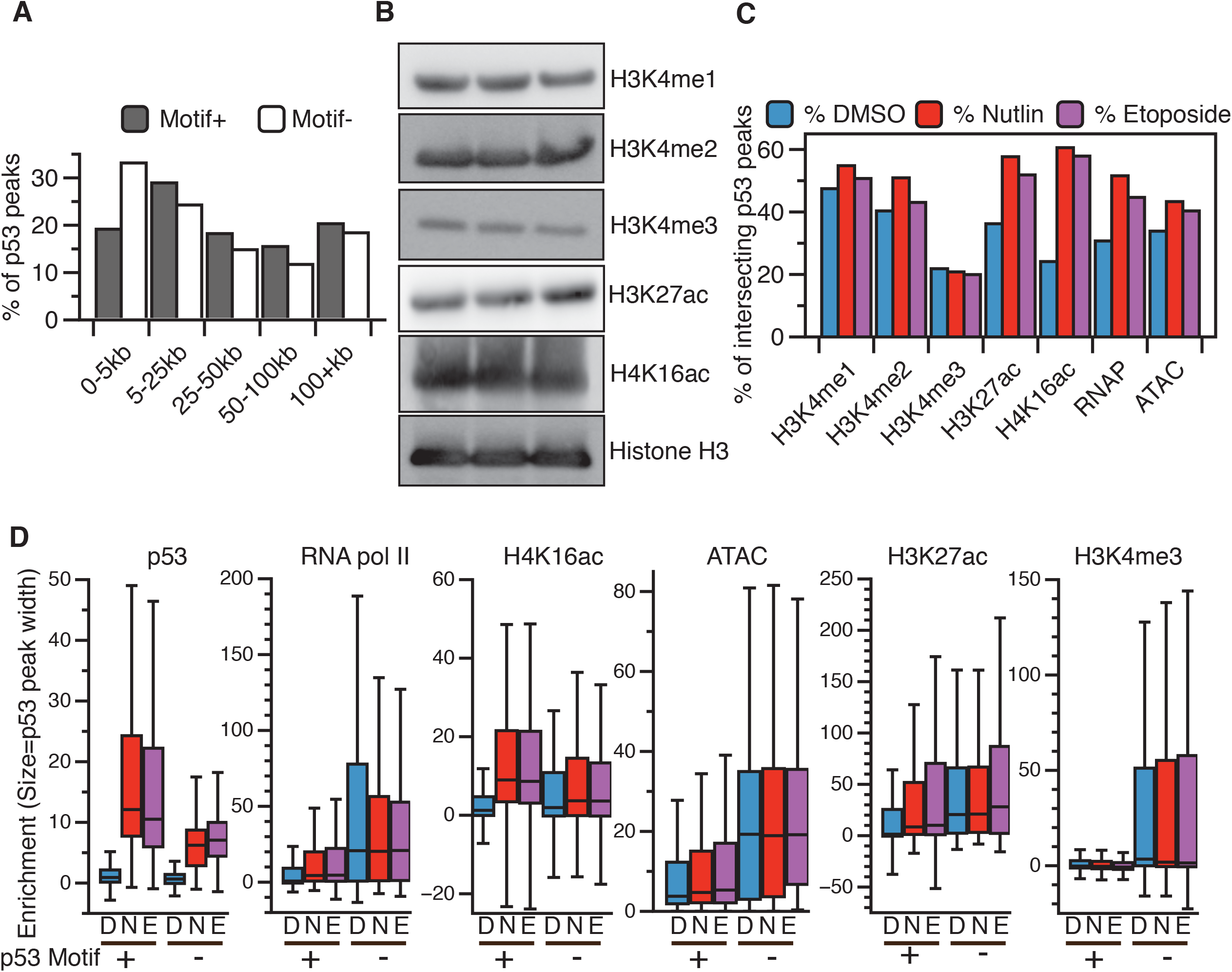
*A*. Western blot analysis of histone modifications used for ChIP-seq analysis from IMR90 fetal lung fibroblasts (cultured at 3% O_2_ in DMEM plus 10% fetal bovine serum) six hours after treatment with either DMSO (vehicle), nutlin 3A (5uM final), or etoposide (100uM final). *B*. Distance of nutlin 3A and etoposide common p53 peaks containing (grey) or not containing (white) a canonical p53 motif from the nearest transcriptional start site (TSS). *C*. Intersection between nutlin 3A and etoposide p53 common peaks and histone modification, RNA polymerse II, or open chromatin (ATAC-seq) peaks as determined by BedTools intersectBed. *D*. Box plot quantification of input-subtracted p53, RNA pol II, H4K16ac, open chromatin/ATAC, H3K27ac, and H3K4me3 across DMSO (D), nutlin 3A (N), and etoposide (E) treatment conditions for motif containing (+) and motif lacking (-) p53 peaks.

Nutlin 3A and etoposide-induced p53 binding events occur within similar local chromatin environments (Fig. 2C). We observe an increase in the number of motif + p53 peaks characterized by *de novo* histone acetylation (both H3K27ac and H4K16ac), increased RNApol II occupancy, and slightly more accessible chromatin (ATAC-seq) after treatment with both nutlin 3A and etoposide (Fig. 2) (34). The local chromatin environment at p53 binding sites are similar between treatments (Fig. 2C), which further supports our previous observations that chromatin structure and modifications are primarily independent of p53 stabilization (27).

We next asked whether there were any distinguishing features of p53 ChIP-seq peaks containing or lacking canonical p53 motifs (2). Motif+ peaks displayed higher input-substracted p53 ChIP enrichment in both nutlin and etoposide conditions compared to p53 motif-peaks (Fig. 2D, p53). This is consistent with previous reports that the p53 motif is the primary determinant for binding affinity (2, 18, 27). Motif-peaks show significantly higher enrichment of RNA polymerase II, H3K4me3, H3K27ac, and ATAC-seq tags than motif+ peaks (Fig. 2D). As motif-peaks are also more closely localized to TSS (Fig. 2A), these data are consistent with technical ChIP-seq artifacts due to actively transcribed and accessible chromatin regions. Treatment-dependent enrichment of RNA pol II, H4K16ac, and H3K27ac relative to DMSO is observed only at motif+ peaks (Fig. 2D), consistent with multiple reports that p53 genome binding leads to the recruitment of RNA polymerase II and transcriptional co-activators like histone acetyltransferase. The H4K16ac-catalyzing enzymes hMOF and TIP60 and H3K27ac-catalyzing enzymes p300/CBP directly interact with and can be recruited to specific genomic loci by p53 (35–38). Taken together, our analysis of local chromatin dynamics reveals strong similarity between p53 binding events downstream of disparate p53 stabilization methods. Further, these data demonstrate ChIP-seq derived p53 peaks lacking the canonical p53 RE localize primarily within accessible chromatin near promoters and are less likely to be observed across p53 activating conditions.

### Transcriptional and promoter dynamics after nutlin 3A- and etoposide-induced p53 activation

Stabilization of p53 via nutlin 3A is highly specific, with very few predicted off-target effects (8). Etoposide, on the other hand, leads to p53 stabilization through failure to repair topoisomerase-induced double-stranded DNA breaks (24). DNA damage itself can activate a number of parallel DNA-damage responsive transcriptional pathways (39). Our data suggest that p53 binding induced by both nutlin and etoposide treatment display similar spatial localization within chromatin, but whether these two treatment conditions produce similar transcriptional responses is not yet known. We therefore investigated whether differential mechanisms of p53 stabilization leads to altered transcriptional activation profiles. PolyA+ RNA from fibroblasts treated with DMSO, 5uM nutlin 3A, or 100uM etoposide for 6 hours was deep sequenced and transcriptome differences between experimental conditions were assessed. Of note, the DMSO and nutlin 3A dataset was previously characterized using identical conditions (27). Using a threshold of 2-fold change between DMSO and the treatment condition, we observe 357 genes upregulated in response to p53 activation downstream of both nutlin and etoposide, whereas 284 genes show reduced expression (Fig. 3A). As expected, commonly upregulated genes include canonical p53 targets involved in cell cycle arrest and apoptosis (Fig. 3B, top)(4). Downregulated genes are strongly enriched in GO categories for cell cycle maintenance and cell division (Fig. 3B, bottom), consistent with a direct role for p53 in transcriptional activation of CDKN1A/p21 and an indirect repression of cell cycle genes through the p21/DREAM complex (40, 41).

**Figure 3.**
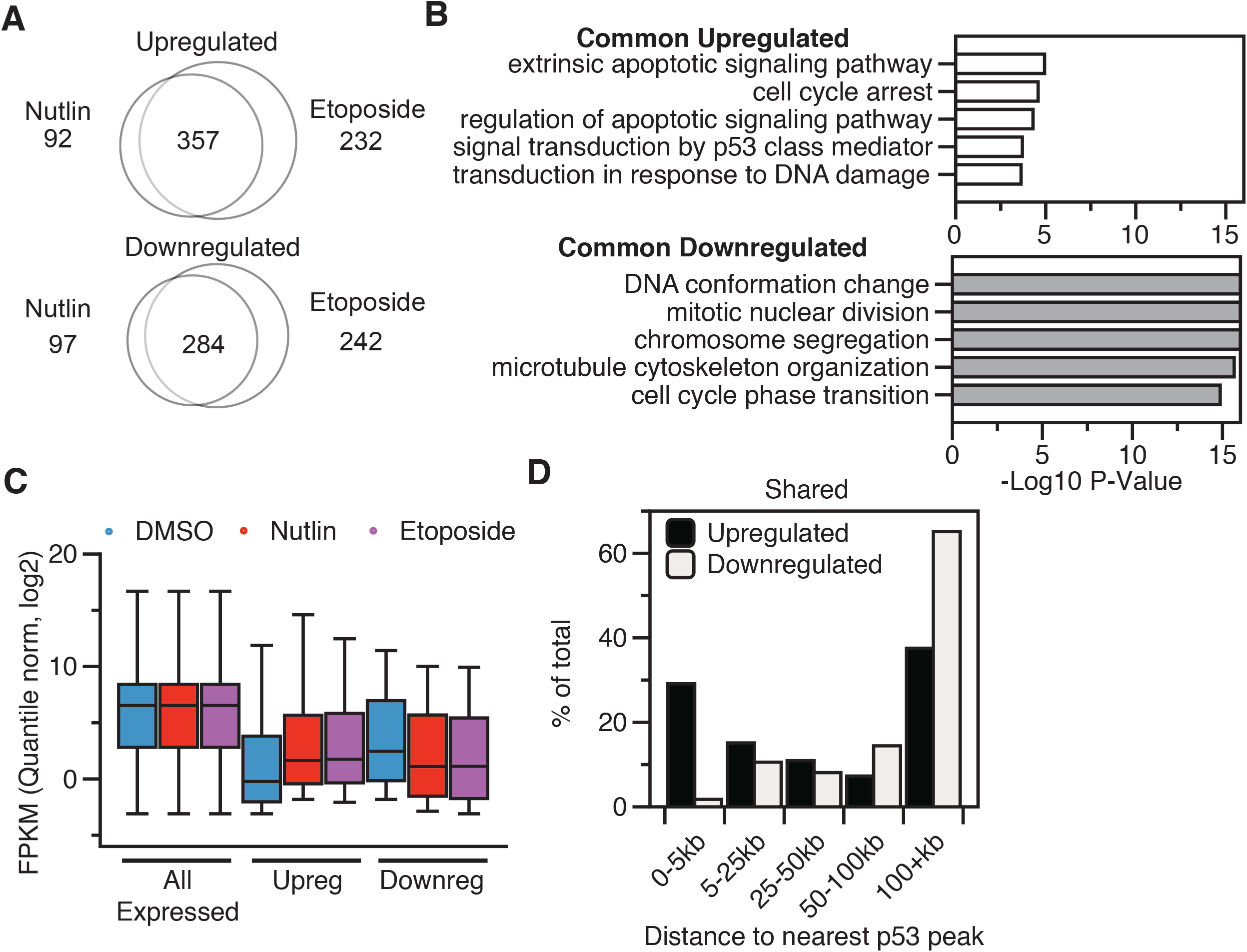
*A*. Intersection between 2-fold upregulated (top) or downregulated (bottom) genes after six hours of nutlin 3A (5uM final) or etoposide (100uM final). *B*. Gene ontology analysis of common upregulated (top) or downregulated (bottom) between nutlin 3A and etoposide treatment of IMR90 fetal lung fibroblasts. *C*. Box plot analysis of quantile normalized FPKM values for all expressed genes (FPKM > 0.1), 2-fold upregulated genes, and 2-fold downregulated genes across DMSO, nutlin 3A, and etoposide treatment conditions. *D*. Distance of upregulated (black) or downregulated (grey) genes to the nearest p53 binding site.

Overall, treatment of IMR90 fetal lung fibroblasts with either nutlin 3A or etoposide yield almost identical transcript expression distributions (Fig. 3C). Nearly 30% of all common upregulated genes have a p53 binding site within 5kB of its transcriptional start site (Fig. 3D), supporting previous observations that proximal p53 binding is required for gene activation (8,23). Of note, over 70% of all commonly upregulated genes are over 5kb from the nearest p53 peak suggesting significant contributions from distal regulatory regions like enhancers. Genes that are commonly downregulated have a skewed distribution, with the modal group of genes displaying p53 binding over 100kb from the nearest gene (Fig. 3D). These results are consistent with the hypothesis that p53 acts solely as a direct transcriptional activator and that downregulated genes are controlled by p53-dependent indirect transcriptional pathways (40).

Measurement of histone post-translational modifications at transcriptional start sites and other regulatory regions has been used extensively to infer transcriptional activity and dynamics (32, 42). Therefore, we assessed changes in chromatin modification status at TSS of p53-responsive genes to discern differences in p53 activating conditions. These analyses also allow the dissection of potential chromatin and transcriptional regulatory mechanisms at p53-activated genes. H3K4me3 and RNA polymerase II, canonical transcriptional start site-associated factors, are enriched at p53 upregulated targets before activation by nutlin 3A or etoposide (Fig. 4A and Fig. 4D). H4K16ac and H3K27ac levels increase at p53-activated target gene TSS after treatment with either nutlin 3A or etoposide (Figs. 4B-C. left). Both of these histone acetylation events are lost at the TSS of genes indirectly downregulated after p53 activation (Figs. 4B-C, right) Downregulated genes have significantly higher transcriptionally associated chromatin modifications and RNA polymerase II occupancy compared to p53-activated target genes (Figs. 4A-D), consistent with the overall higher level of steady state RNA observed for these genes by RNA-seq analysis (Fig. 3C). Pausing analysis (Figs. 4D-E) of p53-activated genes shows increasing RNA pol II occupancy over the gene body of p53 target genes (Fig. 4E, left gene body), but not at the TSS (Fig. 4E, left TSS). This suggests that p53 may influence transcriptional pause release in addition to direct RNA polymerase II recruitment to promoters as has been previously suggested (43, 44). Conversely, downregulated genes display loss of RNA pol II occupancy at both the TSS and along the gene body (Fig. 4E, right) consistent with broad loss of transcriptional activity at these genes in response to p53 activation.

**Figure 4.**
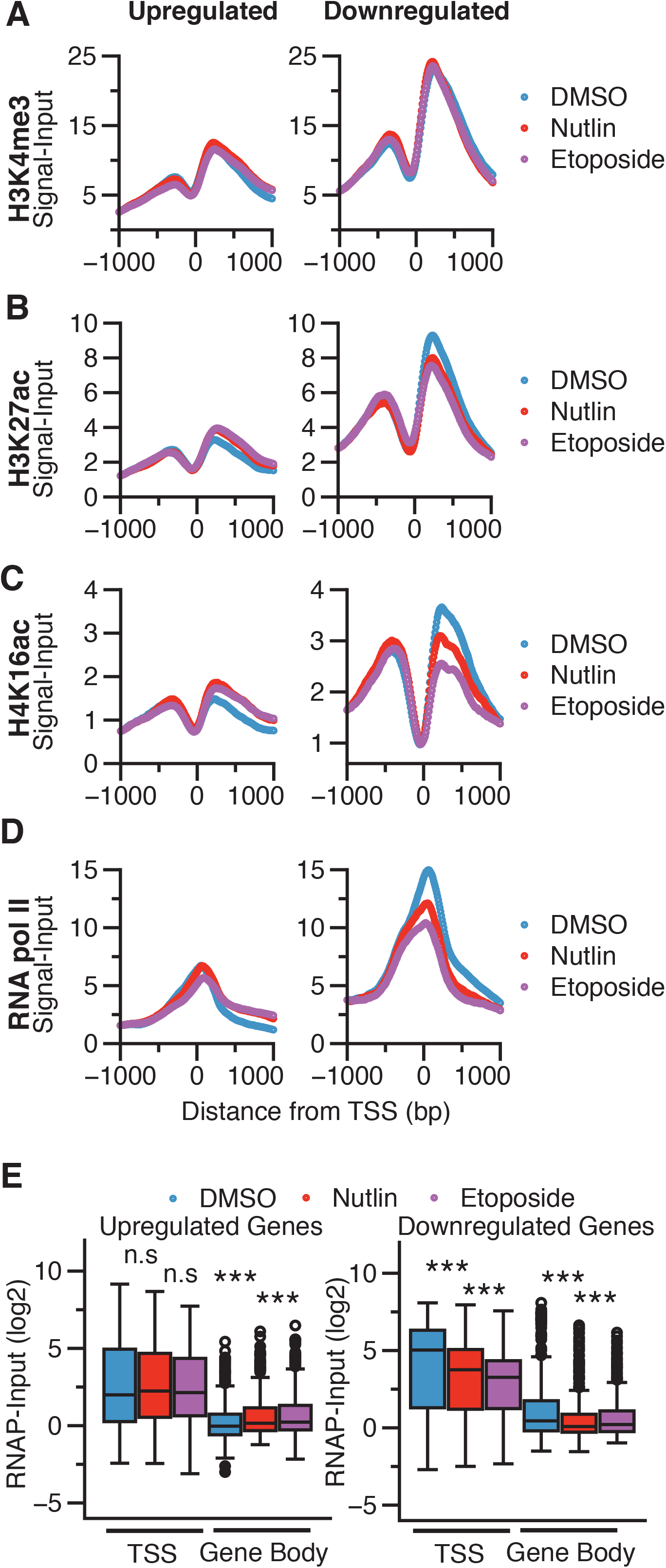
Metaplot analysis at the TSS (-/+ 1000 bp) of 2-fold upregulated (left) or downregulated (right) genes in response to DMSO, nutlin 3A, or etoposide treatment for *A*. H3K4me3, *B*. H3K27ac, *C*. H4K16ac, and *D*. RNA polymerase II. *E*. Box plot analysis of RNA polymerase II occupancy over the TSS or Gene Body of upregulated (left) or downregulated (right) genes. TSS was defined as -/+ 250 bp from the TSS, while gene body was defined as +251 bp to the transcriptional termination site (TTS) of the gene. Significance was defined by using the Mann Whitney *U* test with *** denoting p<0.001.

### Etoposide-specific genes are likely p53-independent, DNA damage-induced NF-kB transcriptional targets

The majority of p53 binding events (Fig. 1D) and induced transcripts (Fig. 3A) are shared between etoposide and nutlin 3A treatment and share similar modes of regulation (Figs. 2D, 4A-E). We next sought to characterize transcriptional differences between our genotoxic and nongenotoxic p53 activating conditions. Less than 100 genes are downregulated or upregulated specifically upon nutlin 3A relative to DMSO treatment (Fig. 3A). These genes fall within three lowly enriched GO categories (Fig. 5A), consistent with previous observations of the high specificity of nutlin 3A for inhibition of MDM2 and subsequent stabilization of p53. Conversely, etoposide treatment induced 232 transcripts 2-fold relative to DMSO treatment that were not found after treatment with nutlin 3A (Fig. 3A). Gene enrichment analysis revealed that these etoposide-specific induced transcripts are related to TNF and inflammatory-dependent signaling (Fig. 5B) (45). DNA damage is a known activator of both p53 and the NF-κB-dependent inflammatory signaling network (39). p53 is implicated in crosstalk with NF-κB in the activation of critical inflammatory genes in immune and epithelial cell types, but not yet in fibroblasts (46–49).

**Figure 5.**
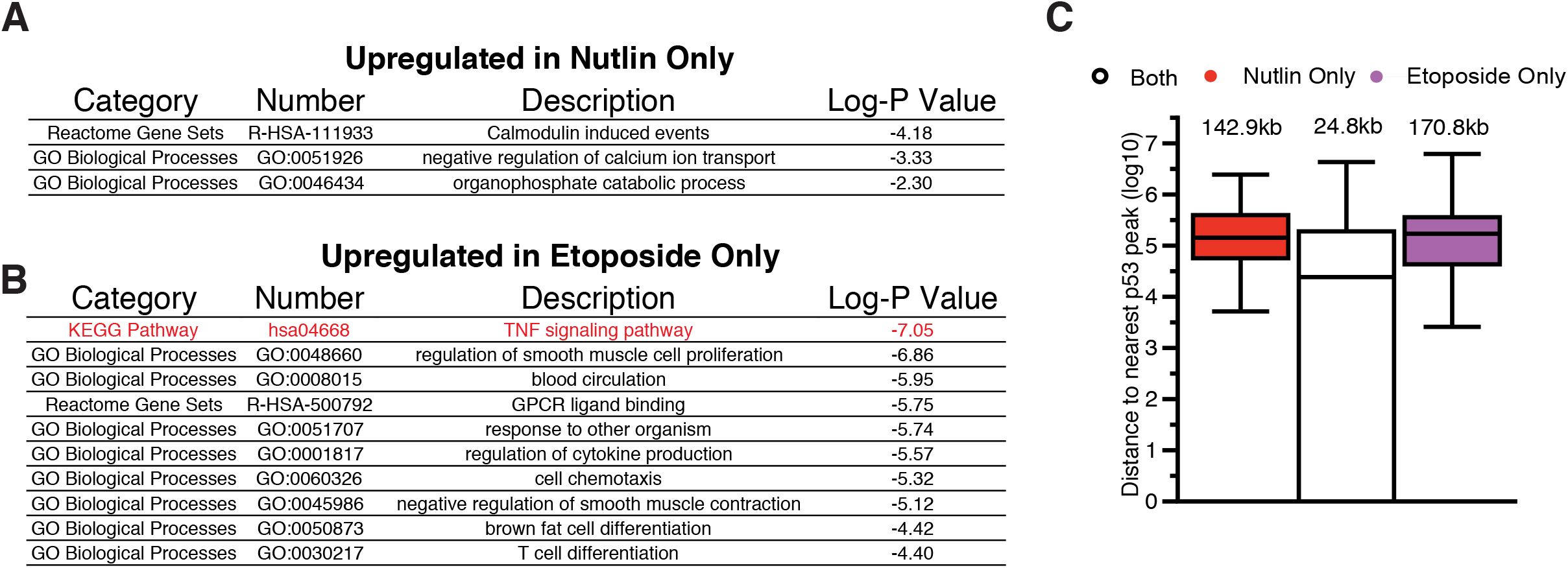
*A*. Gene ontology analysis of 2-fold upregulated genes in nutlin 3A treatment conditions relative to DMSO vehicle control. *B*. Gene ontology analysis of 2-fold upregulated genes in etoposide treatment conditions relative to DMSO vehicle control. Only the top 10 gene ontology terms are depicted and a full list of GO terms are shown in Supplemental Table 3. *C*. Distance of nutlinspecific, etoposide-specific, or commonly upregulated genes to the nearest p53 motif-containing p53 peak.

We therefore investigated wither p53 is directly involved in inflammatory signaling crosstalk in fibroblasts downstream of etoposide treatment. Our nutlin 3A-induced transcriptome does not show a direct p53-dependent activation of inflammatory target genes (Fig. 5A). As p53 binding events occur more proximally to p53-dependent genes than p53-independent genes (Fig. 3D and (23)), we analyzed the distance of etoposide and nutlin 3A-specific genes to p53 binding sites. The median distance between p53 binding events and common p53 target genes is 24.8 kb (Fig. 5C). This distance increases to over 170 kb and 140 kb for etoposide or nutlin 3A-specific genes, respectively, and is, significantly further than the median distance for *bona fide* p53 targets (Fig. 5C).

Multiple reports demonstrate a direct connection between DNA damage-induced inflammatory signaling and the NF-κB pathway (39, 49). Etoposide-specific induced genes are enriched with inflammatory/TNF signaling targets which are under the control of the NF-kB pathway. We therefore tested the possibility that etoposide-specific activated genes are NF-kB-dependent and p53-independent. The p65 subunit of the NF-κB complex is repressed by the activity of IkB and is derepressed by phosphorylation by IκK (50–52). Bay 11-7082 is a small molecule inhibitor of the Iκ kinase family and suppresses NF-κB pathway signaling by maintaining the inactive state of p65 (53). We performed RT-qPCR for three canonical p53 targets and three etoposide-specific inflammatory targets after treatment with IkK inhibitors and activation of p53 by nutlin or etoposide (Figs. 6A-B). The p53 canonical targets CDKN1A, BBC3, and MDM2 are activated in response to both nutlin 3A and etoposide treatment and are unaffected by co-treatment with Bay 11-1043 (Fig. 6A, ratio paired t test). In contrast, IL8, IL1A, and IL1B are all activated specifically after etoposide treatment (Fig. 6B, * p<0.01, ** p<0.001, ratio paired t test), similar to our initial RNA-seq observations (Fig. 3A). Further, co-treatment with Bay 11-7082 abrogates etoposide-induced expression of these genes (Fig. 6B, * p<0.01, ** p<0.001, ratio paired t test) suggesting these genes are downstream of DNA damage-induced NF-kB signaling.

**Figure 6.**
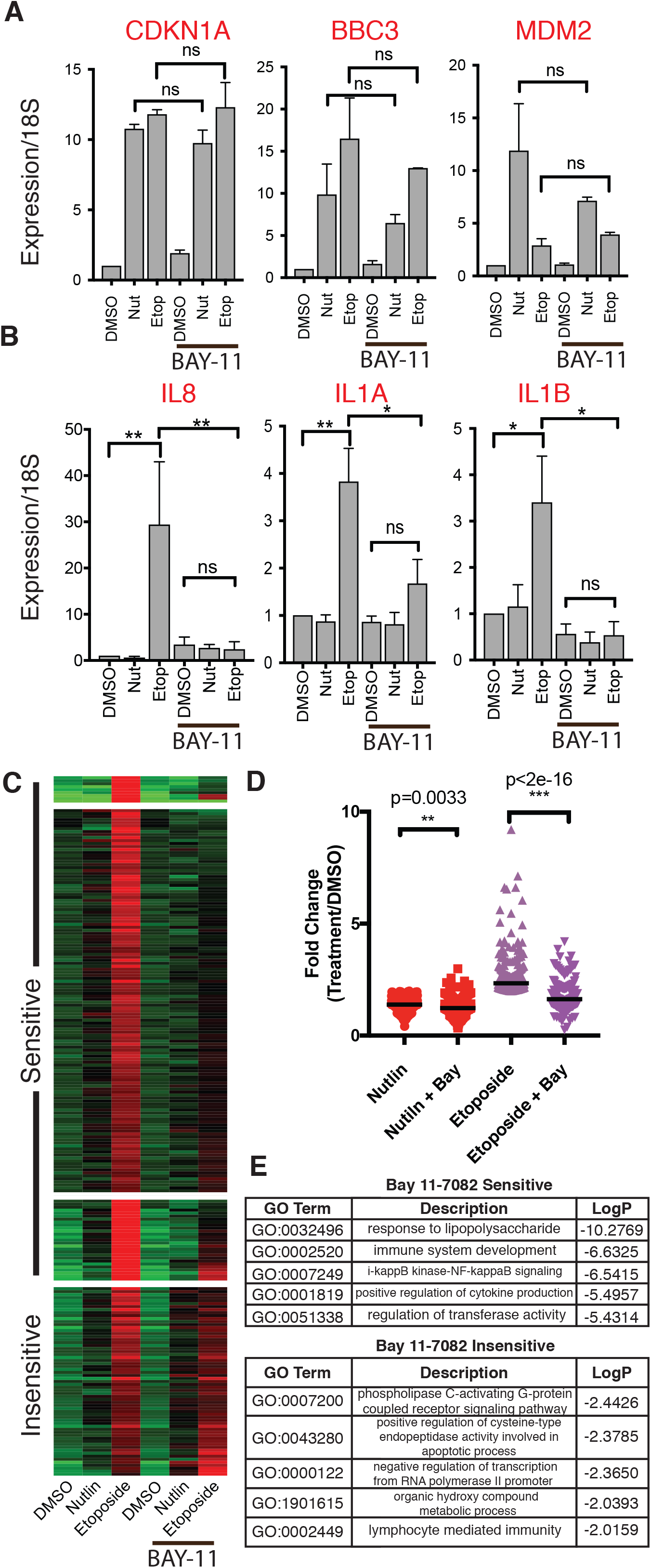
*A*. RT-qPCR analysis of canonical p53 target genes CDKN1A/p21, BBC3/puma, and MDM2 in IMR90 fetal lung fibroblasts in response to treatment with DMSO, nutlin 3A, or etoposide (6 hours total). Samples were co-treated with Bay-11-7082 (5uM) or with additional DMSO. Statistical comparisons were computed using a ratio paired *t* test. *B*. RT-qPCR analysis of etoposide-specific genes CXCL8/IL8, IL1A, and IL1B in IMR90 fetal lung fibroblasts in response to treatment with DMSO, nutlin 3A, or etoposide (6 hours total). Samples were co-treated with Bay-11-7082 (5uM) or with additional DMSO. Statistical comparisons were computed using a ratio paired *t* test. *C*. Heatmap from RNA-seq of IMR90 fetal lung fibroblasts in response to treatment with DMSO, nutlin 3A, or etoposide (6 hours total), with co-treatment with Bay-11-7082 (5uM) or additional DMSO. Data were placed into 4 clusters using *k-*means clustering. *D*. Fold change (treatment/DMSO) of etoposide-induced genes from IMR90 fetal lung fibroblasts with statistics representing results of a paired Mann-Whitney *U* test. *E*. Gene ontology analysis of Bay-11-7082 sensitive (top) and Bay-11-7082 insensitive (bottom) etoposide-specific genes in IMR90 fetal lung fibroblasts. The top 5 categories for each group are depicted and a full list of GO terms are shown in Supplemental Table 3.

We extended this analysis by surveying the polyA+ transcriptome after treatment with p53 activators and NF-κB signaling pathway inhibition with Bay-11-7082. We identified four strong gene clusters in response to co-treatment with p53 activators and Bay 11-7082 treatment using *k-*means clustering (Fig. 6C). Three groups contained genes that are specifically upregulated by etoposide treatment and are strongly repressed when co-treated with Bay 11-7082 (Fig. 6C, Sensitive). Another cluster contained genes whose etoposide-induced expression was insensitive to treatment with Bay 11-7082 (Fig. 6C, Insensitive). Broadly, treatment with Bay-11-7082 reduces expression of etoposide-specific targets after nutlin 3A treatment (Fig. 6D, nutlin vs. nutlin/bay, p=0.0033, Mann Whitney *U)*, suggesting some of these genes are basally regulated by NF-κB signaling. Bay-11-7082 treatment strongly reduces expression of the etoposide-specific gene set relative to no treatment (Fig. 6D. p<2e^-16^, Mann Whitney *U*). Gene Ontology analysis of the Bay-sensitive gene network confirmed these genes are associated with NF-κB pathway and inflammatory signaling, consistent with our hypothesis that these genes are likely NF-κB targets. Genes found in the Bay 11-7082-insensitive cluster were less enriched in total GO terms, but are related to apoptosis and immune signaling. These genes are therefore putative DNA damage-dependent, but likely NF-κB independent, target genes.

## Discussion

Using comparative genomic approaches, we have demonstrated a highly conserved transcriptional and chromatin response to both genotoxic and non-genotoxic p53 stabilization methods. Binding of p53 to chromatin is highly similar across experimental conditions, with the majority of differences attributed to peak calling approaches. This observation is remarkably similar to a recent report of highly conserved p53 binding across cell types and experimental p53 activation methods using a meta-analysis approach (13). One key aspect of this work is the use of a uniform methodology for genome alignment, peak calling, and statistical thresholding across laboratory and experimental conditions. We used a similar approach by first using macs2 to call significant peaks and then creating a combined peak list between experiments (54). Then, we counted tag enrichment within the combined peak regions for both nutlin 3A and etoposide conditions, which yielded strikingly similar enrichment profiles (Figs. 1D-E). Condition-specific peaks (called by macs2) had higher tag counts within the peak region than did the other condition (Fig. 1E), but the overall profile between nutlin 3A and etoposide-induced p53 binding were well correlated (Fig. 1D). Taken together, these data provide additional evidence that p53 engagement with the genome is highly consistent within the same cell type when activated by disparate methods. It is important to note that our analysis was only performed after p53 activation with the MDM2 inhibitor nutlin 3A or topoisomerase II poison, etoposide. Multiple other direct genotoxic activators, such as additional topoisomerase inhibitors, g-irradiation, DNA chemical crosslinking, and UVB-induced pyrimidine dimerization, have been tested for their ability to activate p53-dependent transcriptional signaling, but the genomewide profile of p53 binding has not yet been established for all of these compounds or DNA-damage mechanisms. Further, the binding and activity of p53 downstream of additional p53-activating conditions, like ribosomal stress, reactive oxygen species, nutrient deprivation, or activated oncogenes, are less well understood, opening up critical avenues for in-depth investigation.

DNA damage signaling leads to a number of post-translational modifications (PTM) to p53 (55, 56), especially within the N-terminus (57). These modifications include multiple phosphorylation events in the first transactivation domain of p53, which may help to block the interaction between p53 and MDM2, leading to p53 stabilization. The N-terminus of p53 contains two independent trans-activation domains (TADs), both of which can be extensively modified (57–59). Our data suggest that p53 DNA binding and p53-dependent gene activation are consistent between p53 stabilization methods even though our data suggest that at least serine 15 is differentially phosphorylated between nutlin 3A and etoposide-treated conditions (Fig. 1A). Post-translational modifications to p53 have been directly implicated in differential gene activation and cell fate (35, 36, 51), but their temporal and spatial distribution in the genome is virtually unknown. ChIP-seq of mouse p53 serine 18 phosphorylation closely mirrored results seen with pan-p53 antibodies (60). Mutation of p53 lysine 120 to arginine (K120R) alters p53 genome binding consistent with the predicted role of K120 acetylation in DNA contact (35, 36, 61), but whether the genomewide shift in binding is due to loss of acetylation or altered DNA contacts with arginine has yet to be determined. Ultimately, whether differential p53 stabilization methods yield different patterns of p53 modifications, and whether these directly alter p53 DNA binding, are still open questions. Our data indicate that serine 15 phosphorylation does not drive p53 binding or transcriptional differences in fetal lung fibroblasts, although we note the single six hour-post treatment time point used in our experiments. Additional time points should be examined to determine whether the method of stabilization alters direct p53 activities.

Recent work suggests that individual p53-dependent transcriptional pathways are dispensable for tumor suppression (23), consistent with previous reports that canonical p53-dependent pathways like cell cycle arrest and apoptosis are also not required (62). Our comparative analysis revealed that DNA damage paradigms, in this case with the platinum-based chemotherapy drug etoposide, activate a parallel transcriptional response most likely controlled by the NF-κB transcription factor and not directly by p53. Interestingly, p53 directly activates IL6 and CXCL8/IL8 in primary macrophages (49) and IL1A and IL1B in primary mammary epithelial cells (21). p53 specifically binds to epithelial-specific enhancers upstream of both IL1A and IL1B in mammary epithelial cells (21, 63) but does not bind to these regions in lung fibroblasts (this work) or dermal fibroblasts (21). Here, we find no evidence that p53 directly activates these immune regulatory genes in primary fetal lung fibroblasts, including no change in transcript levels, RNA polymerase occupancy, or p53 binding. Multiple biological and technical differences between experimental conditions may explain the discrepancies between these datasets. First, our primary fibroblasts were cultured under physiological 3% O_2_ conditions which yields lower levels of reactive oxygen species (ROS) than standard 20% O_2_ conditions used in other experimental systems (64). ROS are well-known activators of DNA damage (65) and are involved in significant crosstalk with inflammatory and NF-kB signaling (46, 66). Higher relative levels of ROS may prime p53 towards activation of inflammatory genes in collaboration with NF-κB (49). Alternatively, activation of p53 in cells with high ROS levels could co-activate NF-κB signaling and lead to expression of an inflammatory gene cascade. The underlying mechanisms of cross-talk between p53 and NF-κB, along with other stress-dependent transcriptional networks, represent an important and active area of investigation for both the immunology and cancer biology fields.

A second putative mechanism driving the observed differences in inflammatory gene expression relates to differential p53 activity across cell types. Thus far, inflammatory target gene expression downstream of p53 activation has been studied across varied types of primary and cancer derived cell lines. Every cell type is characterized by a unique collection of active and accessible regulatory elements (42), and p53 binds primarily to active promoters and enhancers (8, 21, 27). The recent comprehensive meta-analyses of the majority of published human p53 ChIP-seq datasets (13) suggests high similarity of p53 binding across cell types. One caveat is that the majority of the analyzed data were from either mesenchymal fibroblast cell lines or transformed cell lines. A conserved core group of p53 binding sites across three cancer cell lines was also recently observed (23), but of note, each cell type had a unique spectrum of binding events. Two recent works suggest that cell type-dependent chromatin accessibility leads to varied p53 binding, which could explain differential p53-induced inflammatory target genes (21, 22). An analysis of 12 transformed human cell lines demonstrates specific p53 binding to cell type-specific accessible chromatin(22), including specific p53 binding to the IL1A locus in the metastatic melanoma LOXIMVI cell line. In primary mammary epithelial cells, p53 binds to two separate active enhancers between IL1A and IL1B and leads to a p53-dependent activation of those genes (21). The chromatin modification and accessibility-based markers suggest these enhancers are inactive in skin or lung fibroblast and that p53 is unable to bind to these regions (21, 27). Our data demonstrate that these genes are not activated by p53 in lung fibroblasts in response to nutlin 3A or etoposide treatment (21, 27). Cell type-specific chromatin accessibility and enhancer activity provides a powerful and intriguing mechanism for differential regulation of p53 target genes.

In summary, our work provides a comprehensive comparison of p53 binding, chromatin state, and transcriptional activity in primary lung fibroblasts exposed to either genotoxic or nongenotoxic activators of p53. We propose that p53 activity and chromatin/RNA polymerase II dynamics are highly correlated within the same cell type regardless of the method of p53 stabilization, and that crosstalk between other DNA damage-activated transcription factors likely contribute to any observed transcriptional differences or cellular phenotypes.

## Materials and Methods

### Cell Culture

IMR90 fetal lung fibroblasts were cultured at 37˚C with 5% CO_2_/3% O_2_ in DMEM (Gibco) with 10% fetal bovine serum and penicillin/streptomycin. Experiments were performed between cell population doublings 20 and 35. Treatments with DMSO, nutlin (5uM final, Calbiochem), etoposide (100uM final) or BAY-7082 (10uM final, Cayman Chemical) were performed for 6 hours before processing cells for downstream experiments.

### Antibodies

Immunoprecipitation and western blotting was performed using the following p53 (Abcam ab80645, clone DO1), histone H3 (Abcam ab1791), H3K4me1 (Abcam, ab8895), H3K4me2 (Millipore, 07-030), H3K4me3 (Abcam, ab8580), H3K27ac (Active Motif, #39133), H4K16ac (Millipore, 07-329), POLR2A (RNA pol II, Santa Cruz, #sc-56767)

### ChIP-seq

Chromatin immunoprecipitation was performed as previously described (27). Briefly, 10 million cells were crosslinked with formaldehyde (1% final concentration) for 10 minutes at room temperature with gentle rotation and quenched with glycine. Cells were isolated, washed 2X with ice cold phosphate-buffered saline, and snap-frozen on dry ice. Chromatin was extracted from isolated nuclei and sheared to 300bp average size using a Diagenode Bioruptor Plus. All reactions were performed overnight at 4°C with rotation. Immunoprecipitated DNA was purified by phenol:chloroform extraction and indexed sequencing libraries were prepared using the NEBNext Ultra DNA Library reagents (New England Biolabs). An Agilent BioAnalyzer was used to determine library sizes and the Invitrogen Qubit fluorimeter was used to quantify library mass. Finally, absolute molarity calculations were determined using the Kapa Library Quantification method and libraries were pooled for sequencing per manufacturer’s recommendations.

All ChIP-seq libraries were run with 100bp single-end reads on an Illumina HiSeq 2000 with the exception of H3K4me2 which was performed with 75bp single-end reads on an Illumina NextSeq 500. Raw FastQ files were aligned to the hg19 reference assembly (downloaded from the Illumina iGenomes repository) using bowtie2 (67) and data were analyzed/visualized using Homer, deepTools, and a local installation of UCSC Genome Browser.

### RNA-seq

DNA-free, total RNA was isolated using RNeasy columns (Qiagen) and 1ug was used to extract polyA+ RNA using magnetic poly(d)T beads (New England Biolabs). Strand-specific RNA libraries were constructed using NEBNext Ultra Directional RNA and BioO NextFlex Rapid reagents. RNA-seq libraries were sequenced with 100bp single end reads on an Illumina HiSeq 2000 (initial comparison of DMSO, nutlin, etoposide) and with 75bp single end reads on an Illumina NextSeq 500 (NF-kB inhibitor experiments). Resulting raw data were aligned to the hg19 assembly using TopHat2/Bowtie2 (68). Differentially expressed genes were those with at least 2-fold difference between the treated condition and the comparable DMSO-treated condition.

### ATAC-seq

Assay for transposase-accessible chromatin (ATAC-seq) was performed as described (34). Briefly, proliferating IMR90 cells were treated with DMSO, nutlin 3A, or etoposide as described above and harvested by centrifugation. 50,000 cells were resuspended in ATAC lysis buffer and incubated on ice for 5 minutes before pelleting at 500 x G for 5 minutes at 4°C. Lysis buffer was then removed and nuclei were immediately resuspended in 50uL of transposase reaction mix (1X TD Buffer, 2.5uL of Nextera Transposase). The transposase reaction was incubated at 37°C for 30 minutes before the reaction was stopped by purification with Qiagen MinElute columns. Transposed DNA fragments were PCR amplified using custom indexing primers before sequencing on the Illumina NextSeq 500.

### Data availability

Datasets found in this manuscript are available without restriction through Gene Expression Omnibus GSE58740 (DMSO and nutlin 3A) and GSE115940 (etoposide).

## Acknowledgements

We thank the University of Pennsylvania Epigenetics Institute and the University at Albany Functional Genomics Core for sequencing support. MAS was supported by start-up funds from the University at Albany and the State of New York and NIH R15 GM128049. SLB was supported by NIH R01 CA078831.

